# Birth timing after the long feeding migration in elephant seals

**DOI:** 10.1101/2020.10.02.324210

**Authors:** Richard Condit, Roxanne S. Beltran, Patrick W. Robinson, Daniel E. Crocker, Daniel P. Costa

## Abstract

Northern elephant seals migrate long distances from feeding grounds to raise pups during a brief period on breeding beaches. Since gestation sets a parturition date months in advance, timing of the arrival must be precise. We used satellite-tracked animals to examine this timing, establishing arrival and birth dates in 106 migrating females and estimating how far they traveled in the days just before birth. Females arrived a mean of 5.5 days prior to birth (range 1-11, sd=1.6), and females arriving later in the breeding season cut that pre-birth interval by 1.8 days relative to early arrivers. There was no correlation between female body condition, nor female age, and the pre-birth interval. The last 15 days prior to birth, animals traveled as far as 1465 km. Those furthest from the colony traveled > 100 km per day, three times faster than animals near the colony at the same time. Despite migrations covering several thousand kilometers while pregnant, female elephant seals were able to time their arrival within 6 days, swimming steadily at high speed if needed. This allows them to maintain a precise annual cycle for many years consecutively.

## Introduction

Reproductive phenology, or the timing of reproduction, is a core feature of life history in animals and plants. A prominent phase of the reproductive schedule in migratory animals is the transition between migration and reproduction. In birds, the phrase ‘arrival biology’ is used to describe this transition, known variously as the pre-chick, post-arrival, pre-breeding, or pre-laying period [[1–5]]. The transition is a small part of the annual cycle, but details of its timing deserve attention. From a distant location, animals must initiate a long migration so that they arrive at the breeding ground on a precise schedule [[6,7]].

Mammals are also migratory, and marine mammals travel long distances between feeding and breeding. Elephant seals (*Mirounga* spp.), for example, spend most of their lives hunting for fish and squid in remote oceans but then migrate thousands of kilometers in order to birth and raise pups on land [[8–11]]. Unlike birds, gestation begins months in advance, setting a parturition date and requiring precise timing of arrival at the breeding colony: too early means wasted foraging time, while too late would be fatal to the newborn. Yet females maintain this precision year after year and give birth on a consistent annual cycle [[12]]. Our broad goal is understanding what limits female reproductive success and how individuals maximize foraging time and pupping success [[13–18]], including the birth timing and whether it limits fecundity.

Here we take advantage of a sample of female northern elephant seals (*M. angustirostris*) that were tracked by satellite during their migration prior to parturition. Satellite-derived locations document movements at the end of the migration as well as the exact arrival time at the colony, while direct observations of pups establish parturition dates. With those observations, we can estimate the time interval between arrival and birth and ask whether 1) more experienced mothers, 2) mothers in poor condition, or 3) late arriving mothers managed to shorten their pre-birth intervals. We also calculated the distance traveled in the last two weeks of the migration to examine females’ ability to control arrival time, asking whether animals further from the colony traveled at a higher speed.

## Materials and Methods

### Breeding and the annual cycle

Northern elephant seals (*Mirounga angustirostris*) breed on remote beaches from Baja California to Vancouver Island. The largest colonies are in central California between 32° to 38° N. latitude [[19,20]]. Females spend 31 days on the colonies in winter where they birth and nurse single pups, then copulate and wean the pup to depart on long foraging migrations (Fig 1). The females are far from shore for 2-3 months, then return to the colony in April or May to molt. At about this time, the fertilized ovum implants in the uterine wall, so gestation takes place during the following seven-month foraging migration in the summer and autumn [[21,22]].

**Figure 1.**
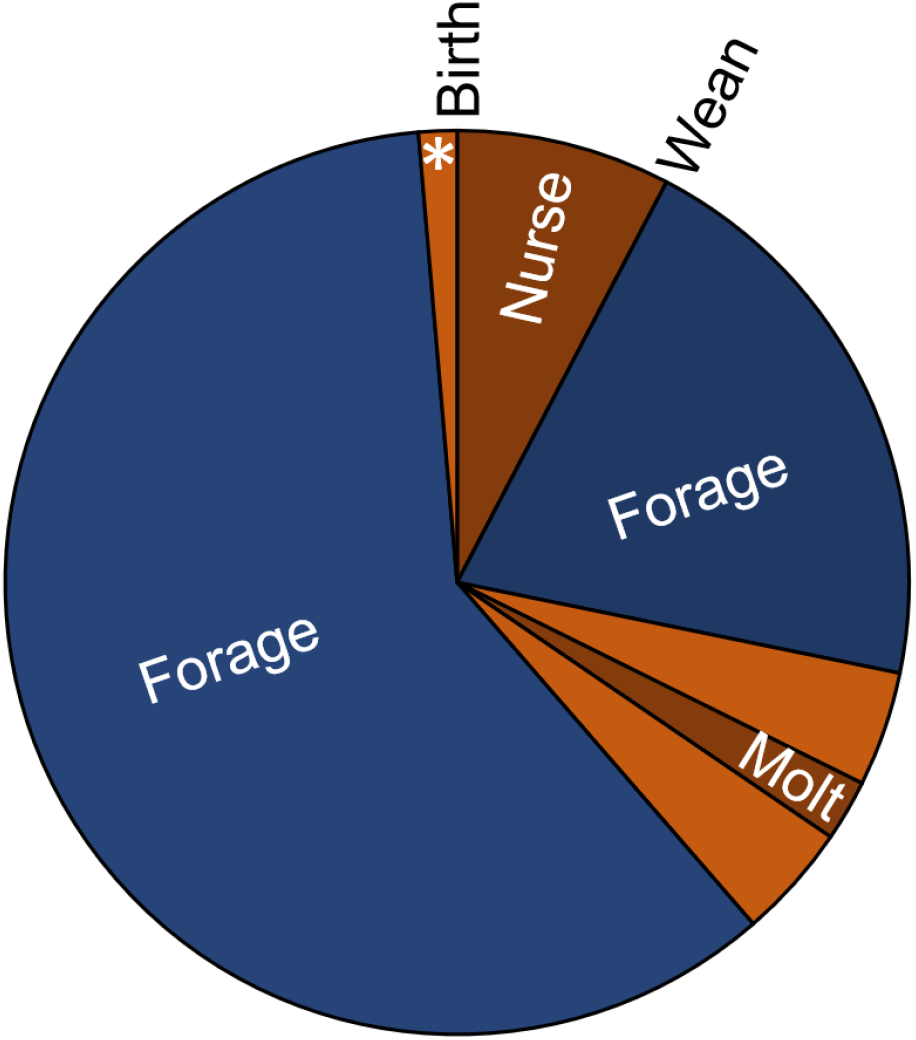
Annual cycle in adult female elephant seals. The year advances clockwise. Periods at sea are in blue, on land in brown. The pre-birth interval is marked with an asterisk.

### Study site and tagging methods

The elephant seal colony at Año Nuevo (37.1086° *N* latitude, 122.3378° longitude) is 31 km north of the University of California, Santa Cruz. Biologists from the university have studied the seals intensively every year since the late 1960s, with a focus on individuals identified by numbered plastic tags attached to their flippers. The age of many adults is known from the tags, allowing individuals to be tracked throughout their 21-year lifespans [[12,23]]. In addition, ARGOS satellite transmitters, GPS, and Time-Depth Recorders were deployed on individuals to document movements while at sea [[8, 9]]. Instruments were attached prior to foraging migrations and removed immediately thereafter [[24]]. While the animals were sedated, detailed morphometric measurements were collected in order to estimate body composition as percent fat [[8,13,25,26]].

### Arrival and parturition

There were 266 females instrumented in May or June between 2002 and 2015, but only 164 were observed on the colony with a pup the following winter, and 30 of those had failed satellite tags or unreliable pup sightings. The remaining 134 records included consistent sightings of the female both before and after birth, but the most precise results come from 106 cases in which there was no gap between the last day seen without a pup and the first with a pup. Ninety-four of those carried a Time-Depth recorder, revealing depth every 8 seconds, revealing arrival time on the beach with high precision. The other 12 females did not have a depth record, only a position from the satellite tag, and we estimated arrival when a high-quality ARGOS location was < 1 km of Año Nuevo and remained there until the animal was observed [[24]]. In all 106 of these cases, the time of birth was observed with a precision of 24 hours, because observations of pups were confined to daylight hours during the short winter days. Though we sometimes see births at Año Nuevo, given 15 hours of winter dark combined with the large number of females on the colony, most are missed, and we did not directly observe the birth in the 106 cases reported here.

### Data analysis

For these 106 breeding records, we found the pre-birth interval *D* as *P* — *A*, where *P* was the first date a pup was observed and *A* the arrival date. To those integers *D* we fitted gamma, Gaussian, and Laplace distributions [[27]], based on the likelihood of the observations assuming a Poisson error. The two parameters of each distribution were estimated with a Metropolis sampler, generating posterior distributions and thus 95% credible intervals.

Using standard linear regression, we tested whether the pre-birth interval *D* correlated with the female’s arrival date *A* (*N* = 106), age (*N* = 85 that had been tagged at birth), and body fat upon arrival (*N* = 104 with morphometrics). The 106 breeding records included 92 different individual females, with 14 individuals having two records. Given so few cases in which one female had multiple records, we did not test for patterns within individuals.

The distance from the colony was calculated 15, 10, and 5 days prior to parturition from the ARGOS satellite locations, providing a female was still at sea. For all 106 females, we calculated the net travel toward the colony between day 15 and day 10, which is the colony-distance on day 15 minus the colony-distance on day 10; it was divided by 5 to express net daily travel.

## Results

### Arrival and the pre-birth interval

Females arrived between 28 Dec and 8 Feb, with a mean of 15 Jan (SD=8.3 d). They gave birth *D* = 1-11 days later, between 3 Jan and 9 Feb (mean 21 Jan, SD=7.9 d). The 1-day and 11-day delays were outliers, however, since every other interval *D* was 3-8 days (Fig 2). The sample mean of *D* was 5.49 d, with a standard deviation of 1.53 d. The Gaussian model fit the distribution of *D* better than gamma and Laplace distributions, though even the Gaussian failed to account for narrow peak at 4-6 days (Fig 2). The Laplace distribution captured the steep peak but had a poorer fit in the tails.

**Figure 2.**
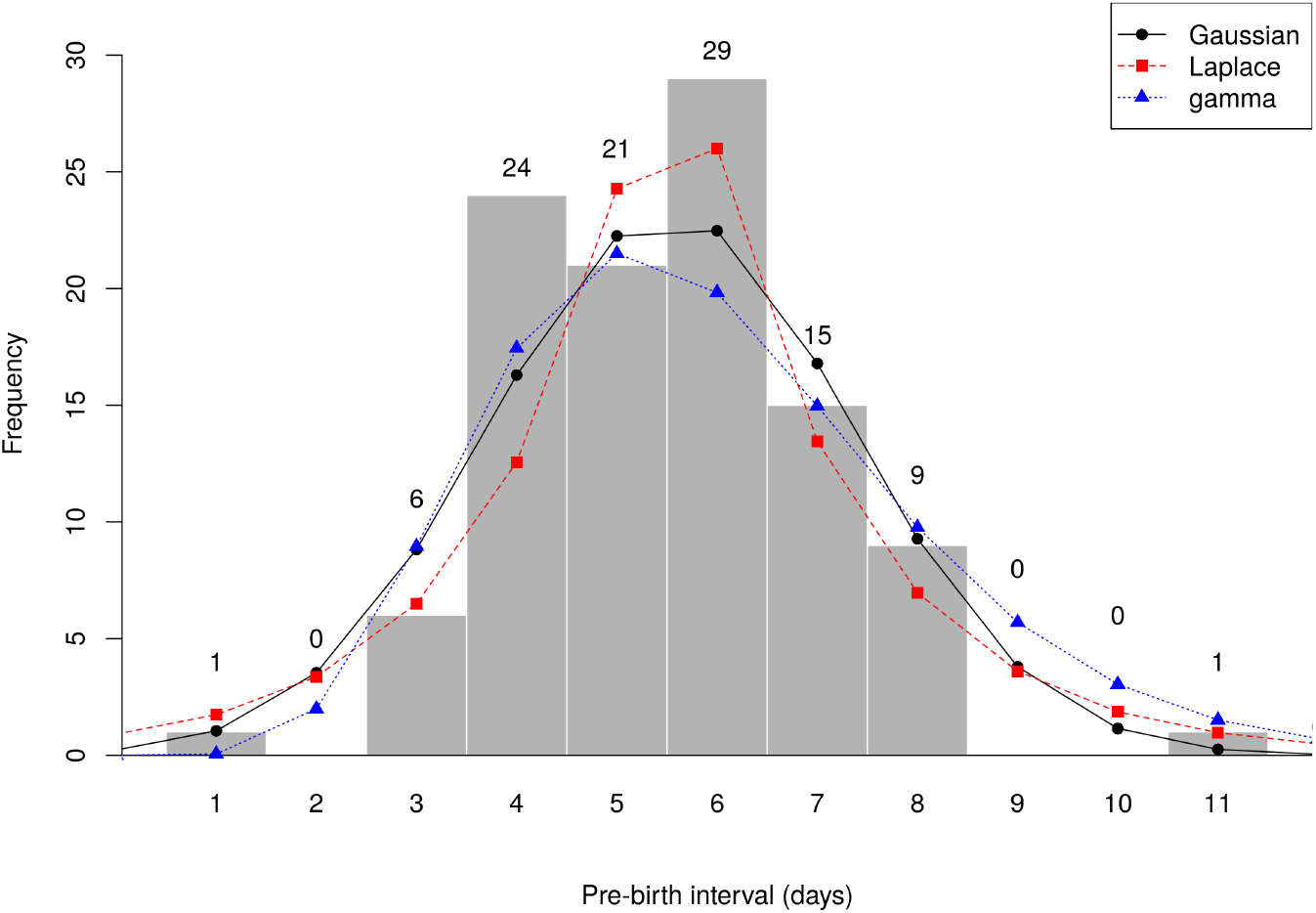
Observed pre-birth interval. Histogram from 106 breeding records (gray bars with numbers above) and fitted distribution from three models. The Gaussian had the highest likelihood (by 1.4 log-likelihood units over the Laplace and 1.6 over the gamma). The best-fit Gaussian had mean = 5.53 (95% credible interval 5.25-5.93) and SD = 1.82 (1.51-2.22).

### Variation in the pre-birth interval

There was a significant negative relationship between arrival date and the pre-birth interval (Fig 3). The mean interval for females arriving on 1 Jan was 6.3 days, decreasing to 4.5 days for females arriving on 1 Feb. There was no relation between female age and the pre-birth period (*r*^2^ = 0.02, *p* = 0.18), nor between a female’s condition (body fat) and the interval (*r*^2^ < 0.01, *p* = 0.64).

**Figure 3.**
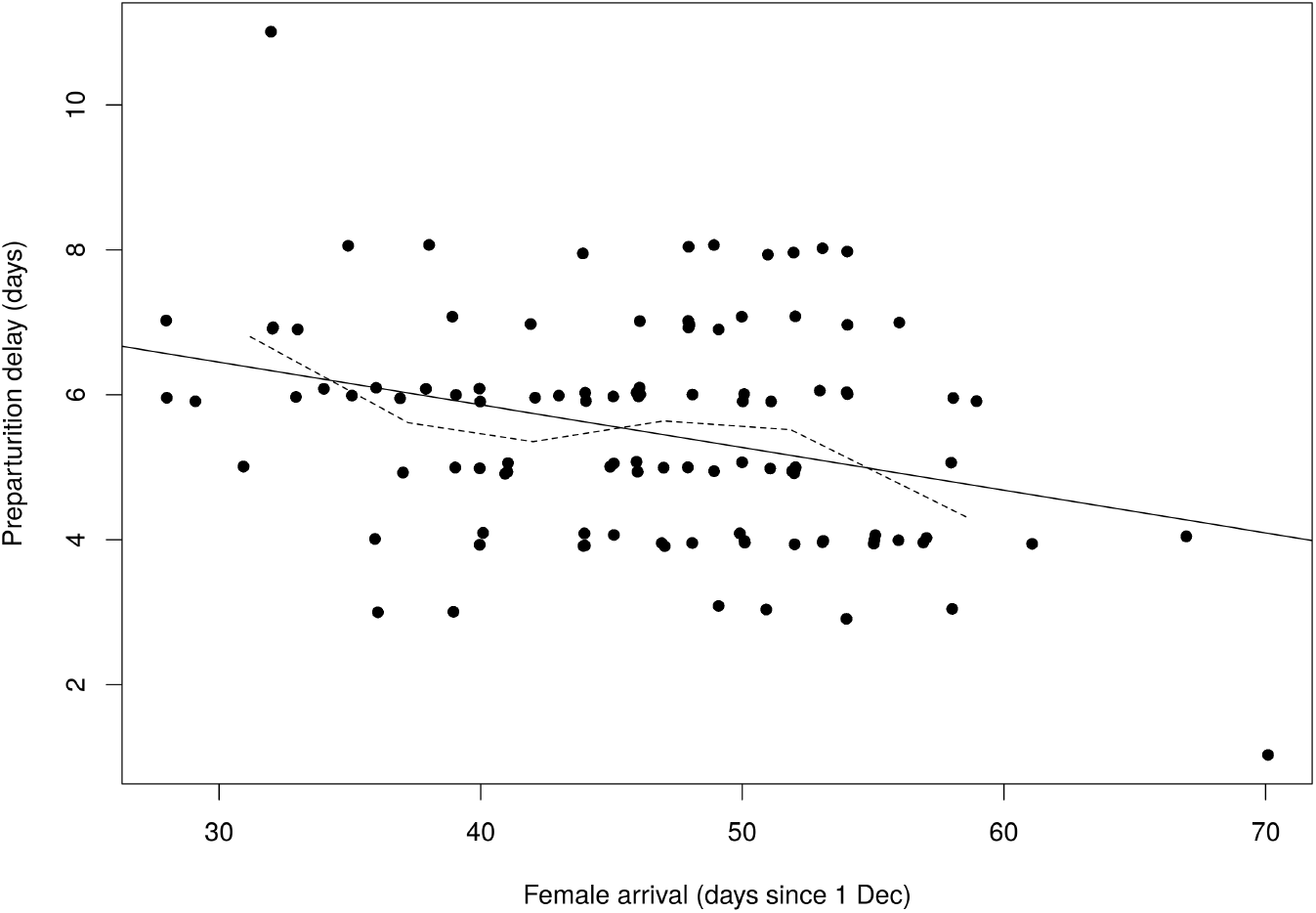
Pre-birth interval versus arrival date. Regression from 106 breeding records. Arrival is given in days since 1 Dec, so day 32 is 1 Jan, day 63 is 1 Feb. The regression coefficient is -.059 (*p* < 0.001, *r*^2^ = 0.10). The solid black line is the regression; the light gray curve connects the mean delay in 5-day intervals (< 35, 35-39, 40-44 … ≥ 55). Points were moved slightly at random to reveal where multiple points coincided.

The shortest pre-birth interval was a single female with a pup one day after arriving (Figs 2, 3). She arrived just after midnight and was seen without a pup on the first day, then with a pup the following day, so the birth happened 15-30 hours after arrival. She was also the latest of all the females to arrive, on 8 Feb, 24 days later than the average. All other females had at least a 3-day delay between arrival and parturition.

The single female who waited 11 days, 3 days longer than any other (Fig 2), was observed 16 times over those initial 11 days, by many different observers, never with a pup. She was then seen only four times with a pup, and the pup was marked during the procedure to retrieve her satellite tag. Subsequently, for her final 20 days on the colony, she was separated from her marked pup. She arrived on 1 Jan, well before average but not the earliest (Fig 3).

### Migration rate prior to birth

Fifteen days prior to parturition, when all females were still at sea, the mean distance from the colony was 681 km; the distribution of distances was nearly symmetric, and the median (650 km) was only slightly lower than the mean. There were 11 females > 1000 km away, and the furthest was 1465 km. At 10 days preparturition, the mean distance was 369 km, with the furthest female 941 km away. At five days prior, nearly half the females were on the colony; the rest were an average of 130 km away, and the furthest 448 km. Between 15 and 10 days before parturition, females traveled an average of 316 km (63 km day^−1^), and there was a strong positive correlation between distance at day 15 and how far they traveled (Fig. 4).

**Figure 4.**
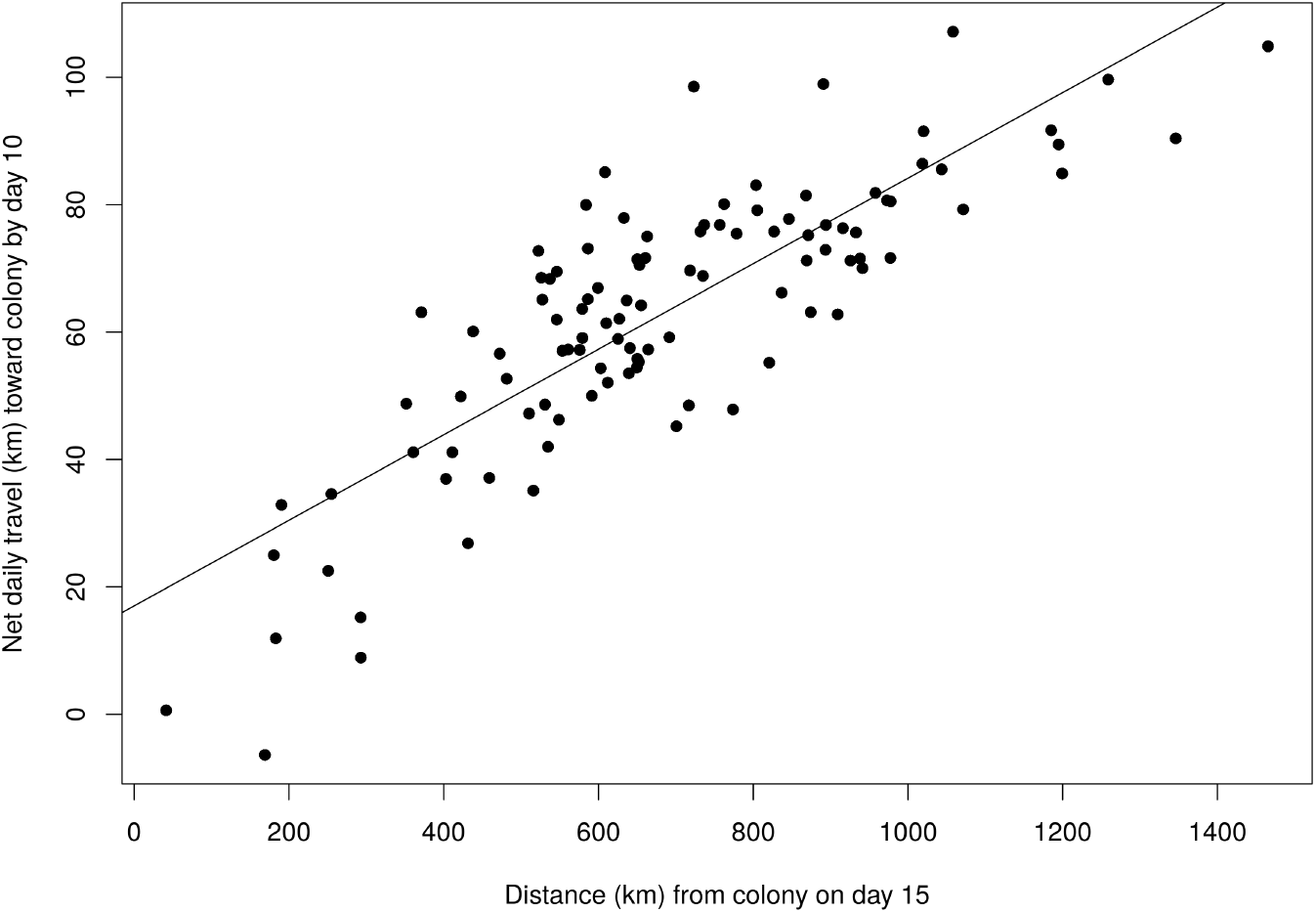
Pre-birth distance traveled. Distance from the colony 15 days before birth (x-axis) versus net travel per day toward colony in the next five days (y-axis). The slope is +0.067 km faster daily travel for every km further from the colony (*r*^2^ = 0.68, *p* < .001, *N* = 106).

## Discussion

Female elephant seals arrived on the colony 5.5 days before giving birth, ranging from 1-11 days. Given how far they traveled over the seven months beforehand and the certain death of a pup born while at sea, this margin did not leave much room for error. Several of the females traveled > 80 km per day in the last two weeks. The observed correlation between speed of travel and distance from the colony suggests the females knew where they were and how soon the pup was due. The fact that most females timed their birth 3-8 days after such a lengthy trip further demonstrates the precision of their cycle.

Selection should favor females who can reduce the delay between arrival at the breeding site and birth, because the delay wastes both foraging time and nursing time. The same selection is expected in other long-distance migrants. In cases where the delay has been quantified, caribou gave birth 4 days after arrival [[28]] and humpback whales 17 days [[29]], while pied flycatchers laid eggs 10 days after arrival, black-throated blue warblers 20 days [[6]], and salmon 20 days [[30]]. White storks, though, waited a much longer 59 days [[31]]. There are important contrasts among these species, however, since gestation occurs throughout the migration in the large mammals, but birds develop eggs after arrival. Shorebirds in Greenland offer a well-studied example of the latter, because the eggs and young are provisioned entirely by local feeding [[3, 32, 33]], whereas in elephant seals and whales, all provisioning – gestation through nursing – is supported by foraging at distant feeding grounds. The birds are thus under selection to adjust the pre-breeding delay in response to food availability on the breeding ground [[4,34,35]], a factor irrelevant in elephant seals and whales. Overall, the short delay in elephant seals and caribou, 4-5 days, can be traced to gestation while migrating. Birds delay laying until 2-3 weeks after arrival because of the conflicting pressures created by short-term egg development and local food supply.

Marine mammals and other large mammals that provision reproduction far from the breeding site are still subject to selection pressures on the timing of birth. Pupping in elephant seals may be timed so that recently weaned pups begin foraging during spring upwelling [[36]]. Another possible constraint is rising temperature on the beach as spring progresses, since warm air temperature is costly to seals [[37]]. Elephant seal females give birth every year, sometimes for many years consecutively [[12]], and they do so on nearly the same date every year, presumably due to these constraints on timing. Our observation that late arriving females cut their pre-parturition interval by nearly two days, and that a single very late female gave birth within 30 hours of arrival, further suggests that females are under pressure to birth within a short winter window.

Assuming that strong selection maintains the birth timing, elephant seal females cannot afford to fall behind from year to year. They must complete the cycle from gestation to nursing inside one year because failing to do so would mean missing a year of reproduction in order to restore the seasonal rhythm [[38]]. Maintaining the annual cycle is thus crucial to a female’s long-term reproductive output [[12]]. Our next goals include elucidating the timing of the full cycle, including reproduction as well as the annual molt (Fig. 1), and how lifetime phenology of individual females affects lifetime reproductive success.

## Conclusion

Female elephant seals maintain a precise annual cycle, giving birth and raising a pup on shore between two long, oceanic migrations. They travel thousands of kilometers late in pregnancy, arriving on shore an average of just 5.5 days before giving birth. Females control their arrival closely, swimming as rapidly as needed during the last two weeks to make that 5.5-day window. They further control the timing of parturition by cutting the delay if they arrive late on the colony. Females are under selection to maintain a tight birth schedule year after year because falling behind would force them to forego reproduction and wait until the following year to raise another pup. They are able to maintain a tight window between the arrival and birth by a precise migration schedule.

## Acknowledgments

We thank B. J. Le Boeuf for initiating and directing the elephant seal research at Año Nuevo for decades, P. Morris and J. Reiter for decades of data collection, and past and present members of the Costa lab, especially field leaders C. Kuhn, S. Simmons, J. Hassrick, M. Fowler, B. McDonald, and S. Peterson. The Institute of Marine Sciences at U.C. Santa Cruz provided funding and logistical support, and California State Parks Ano Nuevo Park rangers provided access to the colony. Support was provided by the US Office of Naval Research (grants N00014-00-1-0880, N00014-03-1-0651, N00014-08-1-1195, N00014-10-1-0356) and the Ocean Partnership Programme (grant N00014-02-1-1012). Field work was authorized under permits 14535, 14636, and 21425 from the National Marine Fisheries Service as well as numerous earlier permits.

